# Long-read whole genome sequencing identifies causal structural variation in a Mendelian disease

**DOI:** 10.1101/090985

**Authors:** Jason D. Merker, Aaron M. Wenger, Tam Sneddon, Megan Grove, Daryl Waggott, Sowmi Utiramerur, Yanli Hou, Christine C. Lambert, Kevin S. Eng, Luke Hickey, Jonas Korlach, James Ford, Euan A. Ashley

**Affiliations:** Department of Pathology, Stanford University, Stanford, California, 94305, USA; Clinical Genomics Service, Stanford Health Care, Stanford, California, 94305, USA; Pacific Biosciences, Menlo Park, California, 94025, USA; Department of Medicine, Stanford University, Stanford, California, 94305, USA; Stanford Center for Inherited Cardiovascular Disease, Stanford University, Stanford, California, 94305, USA; Department of Genetics, Stanford University, Stanford, California, 94305, USA; Stanford Cancer Institute, Stanford, California, 94305, USA

## Abstract

Current clinical genomics assays primarily utilize short-read sequencing (SRS), which offers high throughput, high base accuracy, and low cost per base. SRS has, however, limited ability to evaluate tandem repeats, regions with high [GC] or [AT] content, highly polymorphic regions, highly paralogous regions, and large-scale structural variants. Long-read sequencing (LRS) has complementary strengths and offers a means to discover overlooked genetic variation in patients undiagnosed by SRS. To evaluate LRS, we selected a patient who presented with multiple neoplasia and cardiac myxomata suggestive of Carney complex for whom targeted clinical gene testing and whole genome SRS were negative. Low coverage whole genome LRS was performed on the PacBio Sequel system and structural variants were called, yielding 6,971 deletions and 6,821 insertions > 50bp. Filtering for variants that are absent in an unrelated control and that overlap a coding exon of a disease gene identified three deletions and three insertions. One of these, a heterozygous 2,184 bp deletion, overlaps the first coding exon of *PRKAR1A*, which is implicated in autosomal dominant Carney complex. This variant was confirmed by Sanger sequencing and was classified as pathogenic using standard criteria for the interpretation of sequence variants. This first successful application of whole genome LRS to identify a pathogenic variant suggests that LRS has significant potential to identify disease-causing structural variation. We recommend larger studies to evaluate the diagnostic yield of LRS, and the development of a comprehensive catalog of common human structural variation to support future studies.

---

Short-read sequencing (SRS) methods are currently favored in clinical medicine because of their cost effectiveness and low per-base error rate. However, these methods are limited in their ability to capture the full range of genomic variation.^1^ Areas of low complexity such as repeats and areas of high polymorphism, such as the HLA region, present challenges to SRS and reference-based genome assembly. Indeed, with 100 base pair (bp) read length, fully 5% of the genome cannot be uniquely mapped.^2^ In addition, many diseases are caused by repeats that become increasingly pathogenic in a range beyond the resolution of SRS. Another challenge comes in the form of structural variation, and while SRS has been very successful in the genetic discovery of single nucleotide and small insertion-deletion variation, recent findings suggest we have greatly underestimated the extent and complexity of structural variation in the genome.^3,4^

Long-read sequencing (LRS), typified by PacBio^®^ single molecule, real-time (SMRT^®^) sequencing, offers complementary strengths to SRS. PacBio LRS produces reads of several thousand base pairs with uniform coverage across sequence contexts.^5^ Individual long reads have a lower accuracy (85%) than short reads, but errors are random and are correctable with sufficient coverage, leading to extremely high consensus accuracy.^5,6^ Further, long reads are more accurately mapped to the genome and access regions that are beyond the reach of short reads.^1^ Of particular note, recent PacBio LRS *de novo* human genome assemblies have revealed tens of thousands of structural variants per genome, many times more than previously observed with SRS.^3,7^ These capabilities, together with continuing progress in throughput and cost, have begun to make LRS an option for broader application in human genomics.

Here, we report the use of low coverage whole genome PacBio LRS to secure a diagnosis of Carney complex in a patient unsolved by clinical single gene testing and whole genome SRS. The patient is an Asian/Hispanic male, the product of an uncomplicated term pregnancy who was hospitalized for the first 10 days of life for cardiac and respiratory issues (**Figure 1A**). He remained well until the age of 7 years, when, following the discovery of a heart murmur, he was found to have a left atrial myxoma that was surgically removed. At 10 years, he was noted to have a testicular mass that, at orchiectomy, was found to be a Sertoli-Leydig cell tumor. At 13 years, a pituitary tumor was found and initial conservative management was adopted. Aged 16, he was noted to have both an adrenal microadenoma and recurrence of the cardiac myxomata in the left ventricle and right atrium. Blue naevi were reported. He underwent a second surgical resection of the myxomata with uncomplicated recovery. Aged 18, recurrent cardiac myxomata including a right ventricular and two left ventricular tumors were once again resected and a goretex patch was placed in the right ventricular wall. In the immediate post-operative period, he suffered ventricular tachycardia (VT) and cardiac arrest with spontaneous return of circulation. At this time, a genetics evaluation suggested the possibility of Carney complex but clinical genetic testing (sequencing of *PRKAR1A*) was negative for disease causing variation. At age 19, multiple thyroid nodules were noted on ultrasound, and he was also diagnosed with ACTH-independent Cushing's syndrome, secondary to the adrenal microadenoma. At 21, he underwent trans-sphenoidal resection of the pituitary tumor. At this time, he was found to have recurrent myxomata in the left ventricular outflow tract that have subsequently increased in size (**Figure 1B-C**). To date, these have been treated conservatively with anti-coagulation to reduce the risk of stroke. As of 2016, he is under consideration for heart transplantation and the transplant team judged a molecular diagnosis highly desirable prior to cardiac transplant listing. As a result, whole genome SRS was performed. Genomic DNA was purified, and a library was generated using the Illumina^®^ TruSeq^®^ DNA PCR-Free Library Prep Kit, and genome sequencing was performed using the Illumina HiSeq^®^ 2500 System with paired-end 2 x 100 bp reads to a 36-fold mean depth of coverage. The data analysis and variant curation were performed by the Stanford Medicine Clinical Genomics Service. Single nucleotide variants and small insertions and deletions were identified using MedGAP v2.0, a pipeline based on GATK best practices for data pre-processing and variant discovery with GATK HaplotypeCaller v3.1.1.^8^ This analysis pipeline did not identify any variants that would explain the clinical findings in the patient.

**Figure 1.**
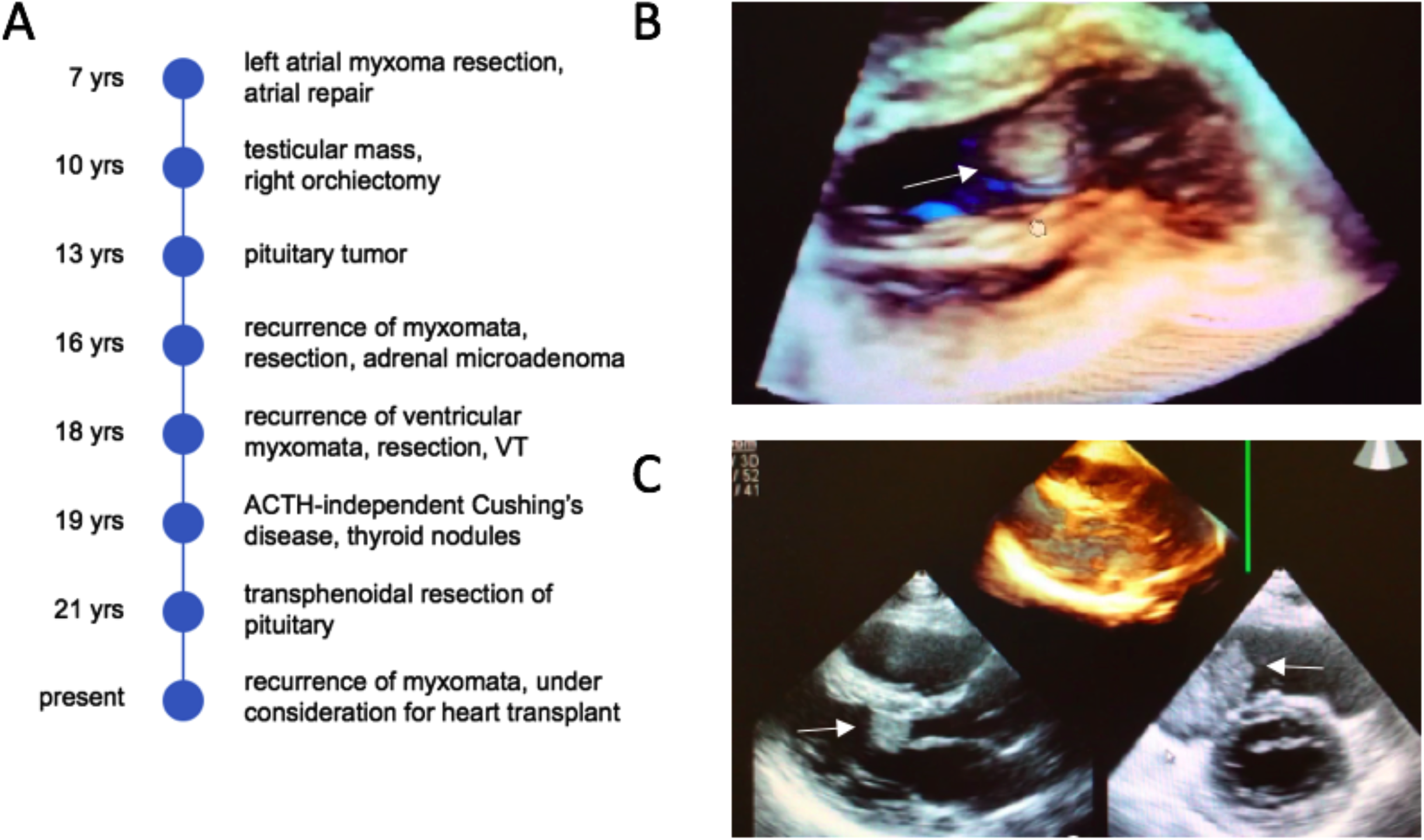
Clinical history and three-dimensional transthoracic echocardiography of patient with multiple neoplasia including cardiac myxomata. (A) Patient narrative. VT= ventricular tachycardia (B) A 2 x 3 cm myxoma is seen in the left ventricular outflow tract (white arrow). (C) The 2 x 3 cm myxoma is seen from another perspective (lower left, white arrow). A 5 x 4 cm myxoma is seen in the right atrium (lower right, white arrow).

To evaluate structural variation, low coverage whole genome LRS was performed on the PacBio Sequel™ system. Following consent under a protocol approved by the Stanford University Institutional Review Board, DNA was isolated from a peripheral blood specimen using the Gentra^®^ Puregene^®^ Blood Kit (Qiagen, Germantown, MD). The DNA was sheared to 20 kb fragments on a Megaruptor^®^ and size-selected to 10 kb using the Sage Science BluePippin™ system. A SMRTbell™ library was prepared and sequenced on 10 Sequel SMRT Cells 1M with chemistry S/P1-C1.2 and 6 hour collections. The sequencing yielded 26.7 Gb (8.6-fold coverage of human genome) in 4.3 million reads with a read length N50 of 9,614 bp. Reads were mapped to the GRCh37/hg19 assembly of the human genome using NGM-LR v0.1.4 with default parameters.^9^ Structural variants were called using PBHoney Spots with ‘-q 10 −m 10 −i 20 −e 1 −E 1 −spanMax 100000 −consensus None’ for deletions and ‘-q 10 −m 70 −i 20 − e 2 −E 2 −spanMax 10000 −consensus None’ for insertions.^10^ Variant calls were further refined to retain only those larger than 50 bp, supported by at least 20% of local reads, and at least 100 bp from an assembly gap.

The resulting call set consisted of 6,971 deletions and 6,821 insertions. To prioritize candidate pathogenic variants, the call set was filtered to exclude variants within a segmental duplication or present in the unrelated control individual NA12878 (A.W., unpublished data). This left 2,368 deletions and 3,174 insertions. Focusing on variants that overlap a RefSeq coding exon resulted in 20 deletions and 16 insertions, with 3 deletions and 3 insertions in genes tied to a genetic disease in OMIM (**Table 1**). Manual review of the 6 candidate variants and correlation with phenotype identified a heterozygous deletion that removes the first coding exon of *PRKAR1A* (NM_212472.2). Germline variants in *PRKAR1A* cause Carney complex, type 1 (MIM #160980), an autosomal dominant multiple neoplasia syndrome.^11^ Two of four reads at the locus unambiguously support the presence of a deletion variant (**Figure 2A**). Because of the random errors in LRS, individual reads from the same allele can have slight disagreements, and two reads can be insufficient to define exact deletion breakpoints with full confidence. Here, the higher quality read supports a 2,184 bp deletion of GRCh37/hg19 chr17:66,510,475-66,512,658 (NC_000017.10:g.66510475_66512658del). This heterozygous deletion variant was validated by Sanger sequencing, which in this case confirmed the precise breakpoints identified by LRS (**Figure 2B**).

**Figure 2.**
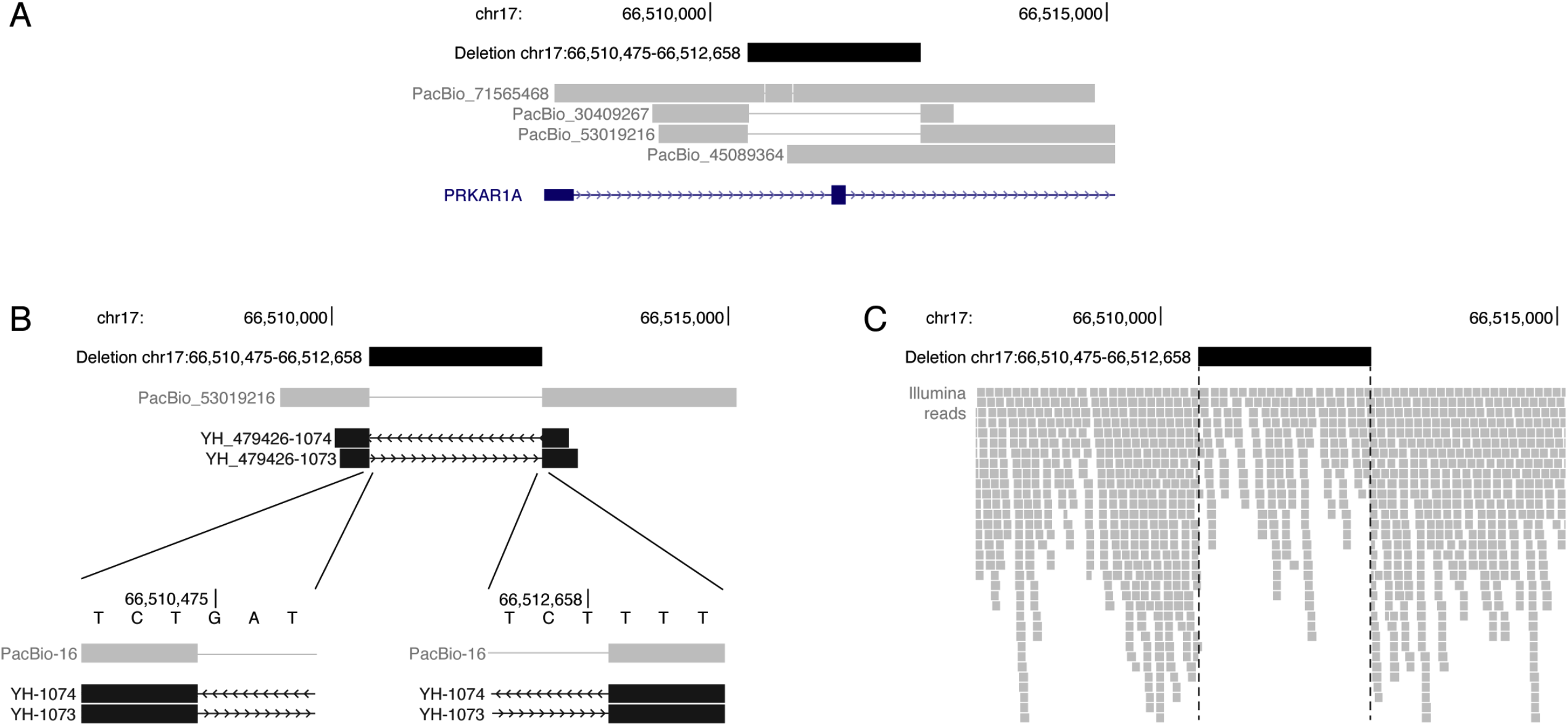
Heterozygous deletion in *PRKAR1A*. (A) PacBio long reads identify a heterozygous 2,184 bp deletion that includes the first coding exon of *PRKAR1A*. Two of four reads at the locus support the deletion. (B) Sanger sequencing confirms the deletion. The forward (YH_479426-1073) and reverse (YH_479426-1074) sequences from a representative amplicon agree to the base pair with the higher quality PacBio read, PacBio_53019216. (C) Illumina short reads support the heterozygous deletion variant through a drop in read coverage and clipped reads at the deletion breakpoints.

**Table 1.** Prioritizing candidate pathogenic variants. The initial call set of 6,971 deletions and 6,821 insertions was filtered to remove variants in segmental duplications or the NA12878 control and to focus on variants that overlap coding exons of genes with a known link to genetic disease.

|  | Deletions > 50 bp | Insertions > 50 bp |
| --- | --- | --- |
| Initial call set | 6,971 | 6,821 |
| Not in segmental duplication | 5,818 | 6,254 |
| Not in NA12878 control | 2,368 | 3,174 |
| Overlaps RefSeq coding exon | 20 | 16 |
| Gene linked to disease in OMIM | 3 | 3 |

It is difficult to call structural variants in SRS data with simultaneously high sensitivity and specificity that is necessary for clinical laboratory testing. Nevertheless, once a small candidate gene list or approximate breakpoints are known, many variants can be identified retrospectively.^5^ In such cases, SRS often provides exact breakpoints to refine the variant discovered by LRS.^12^ Manual inspection of SRS data from the *PRKAR1A* locus shows support for the heterozygous deletion through a drop in read depth and alignment clipping at the deletion breakpoints (**Figure 2C**). Multiple short-read structural variant callers, including Pindel, Lumpy, BreakDancer, Manta, CNVKit, and CNVnator, were retrospectively used to identify structural variants.^13–18^ All tools were run with default parameters. Pindel, Lumpy, BreakDancer, and Manta all identify a deletion in the locus. Pindel and Manta approximately match the breakpoints identified from LRS and Sanger sequencing.

This case demonstrates the ability of whole genome LRS to detect causal structural variation in a rare disease, and to our knowledge, this is the first reported application of whole genome LRS to identify a pathogenic variant in a patient. Although manual inspection of the aligned read data and short-read structural variant callers are able to identify this 2,184 bp deletion, these approaches are not practical to apply genome wide due to limited throughput and high false-positive call rates, respectively. Looking forward, clinical-grade genomics demands strong precision and recall across the full spectrum of genetic variation. SRS has limited sensitivity for variants larger than a few base pairs, and it can miss up to 80% of the structural variants in an individual genome.^3^ LRS appears to be capable of identifying much of the missed variation, and manifests high recall of structural variants even at low depths of coverage.^12^

To accelerate the adoption of sequencing-based structural variant analysis into clinical practice, it will be important for the community to develop and expand an ecosystem of tools and databases similar to that which has arisen around smaller variants. We advocate further development and continued evaluation of tools and best practices for calling structural variants from SRS, LRS and orthogonal data. Additionally, we recommend that the community prioritize creation of a catalog of structural variation derived from these data sources. Databases of common single nucleotide variation, such as ExAC, have proven incredibly valuable.^19^ We expect that a comparable database of structural variants would be similarly valuable and that building the database from LRS would greatly expand current catalogs such as DGV and dbVar.^20,21^

LRS has seen limited adoption in clinical genomics laboratories, in large part due to the per base error rate and cost. Although the individual read error rate requires higher coverage to provide clinical-grade identification of single nucleotide variants, high coverage is not necessarily required for sensitive and specific detection of larger structural variants. Cost effectiveness will ultimately be judged not on cost per base, but on cost per diagnosis. Larger studies on the diagnostic yield of various approaches using LRS will be required to answer the question of the most cost effective technologies for clinical genomics moving forward.

## Acknowledgements

The authors thank the research subject and clinical care teams for their participation in this research study; Chen-Shan (Jason) Chin for helpful discussions; and Primo Baybayan and Matt Boitano for PacBio library preparation and sequencing.

### Web Resources

OMIM, http://www.omim.org/

